# CloVarS: a simulation of single-cell clonal variability

**DOI:** 10.1101/2024.02.22.581631

**Authors:** Juliano L. Faccioni, Julieti H. Buss, Karine R. Begnini, Leonardo G. Brunnet, Manuel M. Oliveira, Guido Lenz

## Abstract

High-throughput time-lapse microscopy has allowed researchers to monitor individual cells as they grow into colonies and react to treatments, but a deeper understanding of the data obtained after image analysis is still lacking. This is partly due to the biological and computational challenges related to long-running experiments and single-cell tracking. In this work, we present the Clonal Variability Simulator (CloVarS), a Python tool for generating synthetic data of single-cell lineage trees to model time-lapse microscopy experiments. After colony initialization, each individual cell is simulated for a given number of simulation frames. Over the course of the simulation, cells are able to migrate, enter mitosis (divide), and enter apoptosis (die). These events are determined by distributions of cell division and death times, which can be inferred from and fit to experimental data. Colonies have an adjustable fitness memory, meaning cells are able to inherit their parent cell’s fitness (i.e. their parent’s time to division/death) to a higher or lesser degree. Arbitrary treatments are able to be delivered to colonies at any time point, modifying their division and death distributions, and fitness memory. CloVarS is an important asset for quickly exploring colony fitness dynamics, testing biological hypotheses, bench-marking cell tracking algorithms, and ultimately improving our understanding of single-cell lineage data.

## Introduction

A human cell can be understood as a system of biochemical signals that collectively direct it towards a given phenotypic state - growth, migration, division, homeostasis, death - over time. Advances in time-lapse microscopy have allowed researchers to glimpse into these complex relationships through single-cell tracking (1–4), but this is not without its challenges. Maintaining cell viability during long-running experiments may be difficult or nearly impossible, depending on the experimental settings (5). While much effort has been put into understanding single-cell tracking data in many different contexts, currently there is no methodological consensus on how to best analyze them (6). Additionally, the variability generated in clonal cells can stem from different sources and modeling how clones are generated is a key strategy for understanding the mechanisms underlying the onset of clonal cell heterogeneity (7–9).

Here, we introduce the **Clo**nal **Var**iability **S**imulator (CloVarS), a simulator of the formation of cell colonies. The parameters governing cell division and death can be derived from experimental data of treated and untreated cells, thus producing large amounts of biologically relevant data in seconds instead of weeks. CloVarS can be used to test hypotheses of cell growth dynamics and explore novel methods for analyzing single-cell tracking data without the need for a complex experimental setup. CloVarS is written in Python. Its source code, along with installation and execution instructions, can be found at https://github.com/jfaccioni/clovars.

## Materials and methods

We briefly explain CloVarS, with further details provided in the Supplementary Data.

### Simulation model

CloVarS simulates individual clonal cells over time. At each simulation time step (referred to as a *frame*), each cell is able to either: migrate to a new position; divide into two new cells; or die and vanish from the colony (Fig. 1A). At every frame, data from the current simulation state is written to the output files. The simulation ends when a stop condition is met (Supplementary Table S1). Output files can then be further processed or analyzed as desired. For convenience, CloVarS includes some basic visualization and analysis scripts of the output files.

**Fig. 1.**
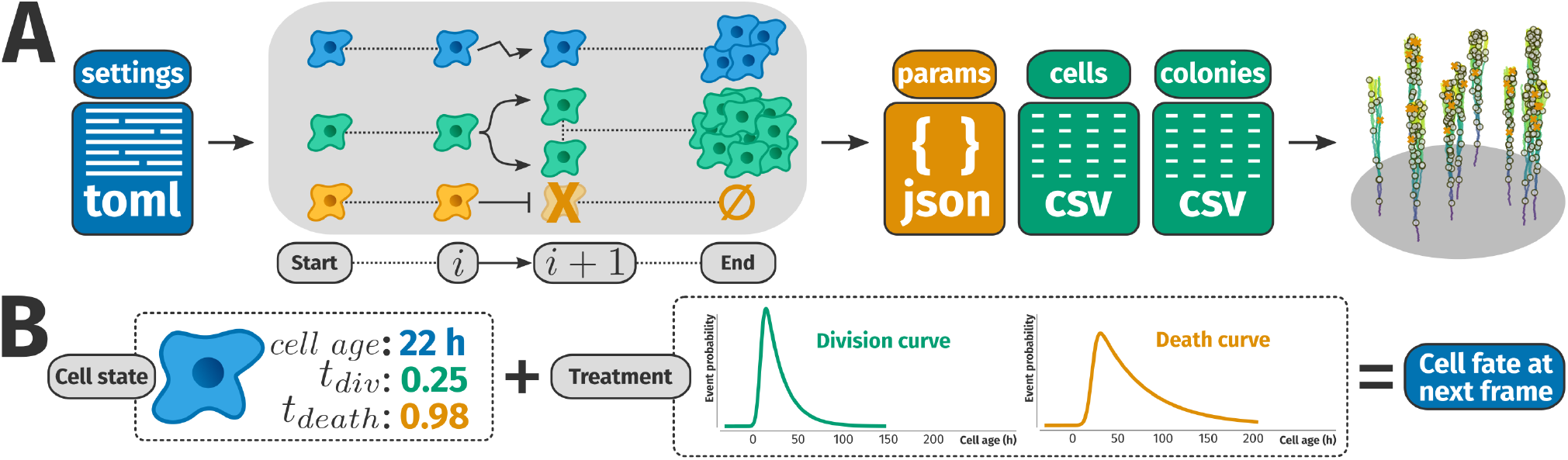
CloVarS overview. (A) The simulation starts after reading the parameters from the settings file (see Supplementary Tables S1, S2, S3, S4 for parameter details). Between adjacent simulation frames *i* and *i* + 1, cells are able to migrate, divide, or die. Once the simulation ends, the output files can be used for data visualization and analysis. (B) Cell fate at the next frame is determined by factoring in its age, division and death thresholds, and the treatment it is subject to (see also Supplementary Fig. S1).

### Fitness thresholds and cell fate

Each cell has a fitness threshold for division (*t*_*div*_) and death (*t*_*death*_) (Fig. 1B). The fitness thresholds indicate how early a cell will trigger its division/death event; for example, if *t*_*div*_ = 0.1, then the cell divides at the age corresponding to the 0.1 percentile of the division curve (see the numerical examples in Supplementary Fig. S1). If the cell has not reached the age for neither of its division or death events to trigger, its fate at the next frame is to migrate. Collectively, *t*_*div*_ and *t*_*death*_ represent the cell fitness, with high-fitness cells having low *t*_*div*_ (early occurrence of division events) and high *t*_*death*_ (resistance to death events).

### Fitness memory and inheritance

At the simulation start, each cell draws its fitness thresholds from a uniform distribution. Upon cell division, these values may be partially or completely inherited by its daughter cells. This is determined by the fitness memory (*f*_*m*_) of the colony: daughter cells from high *f*_*m*_ mothers tend to have almost identical *t*_*div*_ and *t*_*death*_, both amongst themselves and their predecessor. On the other hand, *f*_*m*_ values close to 0.0 means that *t*_*div*_ and *t*_*death*_ values are largely uncorrelated between mother and daughter cells (Fig. 2A). Intermediate *f*_*m*_ values proportionally guide the colony towards fitness preservation (*f*_*m*_ ≈ 1.0) or randomization (*f*_*m*_ ≈ 0.0). *f*_*m*_ is constant and equal for all cells in a colony, although a treatment is able to modify it.

**Fig. 2.**
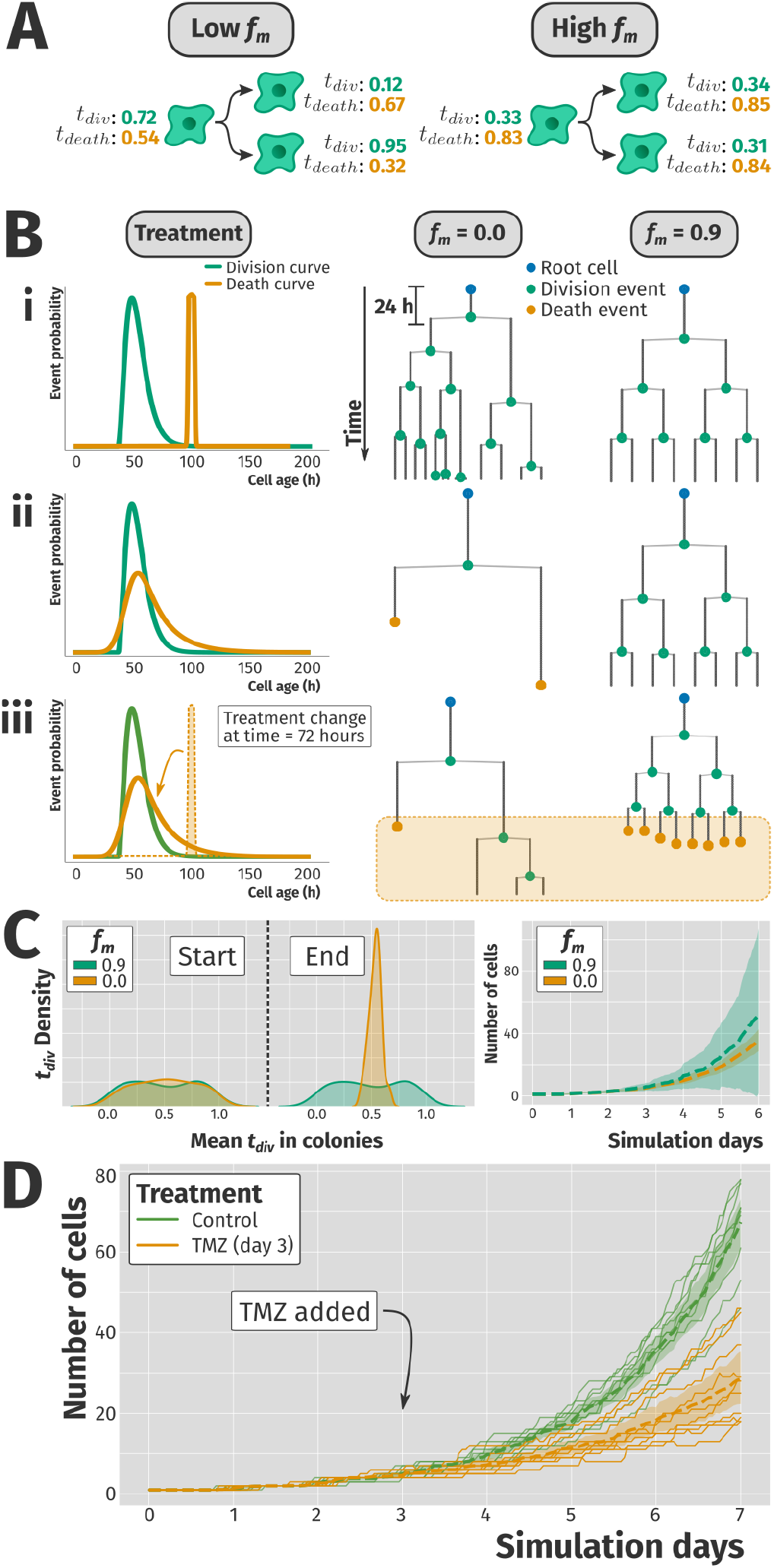
CloVarS results. (**A**) Mother cells with high fitness memory (*f*_*m*_) produce offspring with similar fitness thresholds, while daughter cells with low *f*_*m*_ have largely uncorrelated fitness thresholds among themselves and their mother. Two colonies with a single initial cell each were simulated for 120 frames in the following scenarios: (*i*) no cell death; (*ii*) moderate cell death; (*iii*) moderate cell death after treatment change at 72 h. (**C**) Effect of *f*_*m*_ on colony size variability. For each *f*_*m*_, 100 colonies were simulated for 144 frames. Left, distribution of mean *t*_*div*_ in colonies at the start and end of the simulation. Right, average number of cells + SD (shaded area) for each *f*_*m*_ over time. (**D**) Simulation results based on experimental data. For each treatment, 10 colonies with *f*_*m*_ = 0 were simulated for 168 frames. Dashed lines: treatment average. In Figs. 2B, 2C, and 2D, Δ*t* between frames was 1 h. Code used to produce Figs. 2B, 2C, and 2D can be found in the CloVarS repository.

### Treatments

During the simulation, cells are under a given treatment (the term “treatment” is also used to refer to untreated/control scenarios). Treatments are used alongside an individual cell’s *t*_*div*_ and *t*_*death*_ when defining its fate at the next frame (Fig. 1B). A treatment holds two probability density functions representing a division curve and a death curve, respectively. At population level, these curves describe the chance a cell has to divide or die as a function of its age. The division/death curves for a given treatment can be inferred from experimental data of cell age at division/death, respectively (see Supplementary Data for details). The current treatment can be modified during the course of the simulation.

## Results

### Simulation scenarios

The scenarios presented in Fig. 2B demonstrate the effect of treatments and *f*_*m*_ on colony growth patterns. In scenario *i*, no death events were triggered due to the death curve being dramatically shifted to the right. Cells from the low *f*_*m*_ colony divide at diverse ages, with no correlation between mother and daughter age at division. On the other hand, the offspring of the high *f*_*m*_ colony divides in regular intervals, mimicking the fitness of its initial cell (Fig. 2B, scenario *i*).

When cell death events are introduced, either from the simulation start (scenario *ii*) or after a treatment change (scenario *iii*), having low *f*_*m*_ increases the odds that at least some cells from the colony are able to survive the treatment. This is known as fractional killing in an *in vitro* experimental setting (10). In contrast, a colony with high *f*_*m*_ faces an all-or-nothing scenario: as long as its initial cell is able to survive, its fitness thresholds are largely copied to its offspring, giving rise to a stably resistant colony (Fig. 2B, scenarios *ii* and *iii*). Interestingly, even though low *f*_*m*_ leads to a higher heterogeneity in lineage tree structure, high *f*_*m*_ produces a higher variability in colony sizes (Fig. 2C). In fact, huge colonies are only achievable by high *f*_*m*_ colonies that preserve their fast division time and resistant phenotype throughout generations (Supplementary Video S1), as suggested by experimental data (9).

### Experimental comparison

To evaluate how CloVarS compares to a traditional *in vitro* experiment, we fit the division and death curves from experimental data of manually tracked cells (see Supplementary Data for details). Two treatments were defined: *Control*, based on division times of A172 glioblastoma cells under unhindered growth, and *TMZ*, estimated from division and death times of A172 glioblastoma cells under concentrations of temozolomide that induce fractional killing. Similarly to *in vitro* experiments, cells under the *TMZ* condition were left to grow for 3 days before treatment addition. Switching to the *TMZ* treatment reproduces the fractional killing effects observed *in vitro* (Fig. 2D, Supplementary Fig. S2).

### Closing remarks

Producing accurate and reliable single-cell microscopy data is experimentally and computationally challenging. In this sense, CloVarS can be a valuable tool for generating data for a virtually infinite number of colonies, and testing a variety of biological hypotheses. Cell signaling fluctuations can also be explored and modeled using CloVarS (Supplementary Fig. S3, Supplementary Video S2).

While other biological cell simulators exist in the literature (11–16), a major focus during CloVarS’ development was to keep a biology-first mindset: the underlying rules of the simulation should not be overly complex to the point where the concepts cannot be understood by most cell biologists. The interdependence of cell fitness thresholds and treatment curves when choosing cell fate is a deliberate design choice for the simulation. It establishes a link between the individual cell fitness (*t*_*div*_ and *t*_*death*_) and the population-derived division and death probability density functions. This is in accordance to experimental results that indicate that fitness is a dynamic phenotype that partially depends on the cell itself, but also on its surrounding environment (9, 17).

The interplay among individual cell fitness, colony fitness memory, and treatments are assembled into a program that is stochastic at its core, but reproduces the expected clonogenic behavior and variability as observed *in vitro* (18). CloVarS allow researchers to explore biological hypotheses related to the formation of heterogeneous cell colonies and their fitness dynamics.

## Software availability

An overview of CloVarS, as well as installation instructions, can be found at: www.ufrgs.br/labsinal/clovars.

The source code for CloVarS is deposited at: https://github.com/jfaccioni/clovars.

