## Supplementary Data for "CloVarS: a simulation of single-cell clonal variability"

##### This PDF file includes:

Supplementary Text
Supplementary Figs. S1 to S3
Supplementary Tables S1 to S4
Legends for Supplementary Videos S1 to S2

##### Other supplementary materials for this manuscript include the following:

Supplementary Videos S1 to S2

### Supplementary Text

#### Simulation details

**Overview.** CloVarS utilizes an object hierarchy for modeling its simulation actors: each simulation starts with a **well**, representing a physical well in a cell culture dish. A well is able to hold one or more **colonies**, with each colony being composed of one or more **cells**. Each colony is associated to a **treatment regimen** that describes which **treatment** is currently in effect, and at which frames it is replaced by another treatment (if any).

When CloVarS starts, colonies are created and their corresponding simulation treatments are initialized according to some description found in the configuration file. A well is instantiated, and its initial colonies are placed inside it. The simulation starts at time $t_0 = 0$ seconds, and advances $\Delta t$ seconds at each frame. While the simulation allows for an arbitrary $\Delta t$, users should adjust the value so that it remains lower than the typical time to cell division or death. Empirically, we found $\Delta t = 3600$ (i.e. 1 hour) to strike a good balance between simulation duration and experimental accuracy.

**Simulation frames.** At each simulation frame $f_i$, the following events take place:

- Each colony checks if its current treatment needs to be modified;
- The fate of each cell at frame $f_{i+1}$ is set;
- The simulation state at frame $f_i$ is written to the output files;
- The simulation checks for stop conditions:
  - if at least one stop condition is met, the simulation ends;
  - otherwise, each cell advances $\Delta t$ seconds, triggering their fate at frame $f_{i+1}$.

These steps roughly translate to the following pseudocode:

```
for colony in well:
    colony.modify_treatment_if_needed(delta_t)
    for cell in colony:
        cell.set_fate_at_next_frame(delta_t)  # cell fate is defined here

write_simulation_status_to_output()

should_stop = check_stop_conditions()
if should_stop is True:
    end_simulation()

for colony in well:
    for cell in colony:
        cell.advance_time(delta_t)  # cell fate is triggered here
```

**Advancing cells in time.** When cells advance in time, they trigger a division, death, or migration event. Both death and division events remove the cell from the colony, but a division event also creates two new cells and adds them to the colony. A migration event simply places the cell at a nearby position (randomly determined by Brownian motion) and increases its age.

When a new cell is created, its fitness thresholds for division ($t_{div}$) and death ($t_{death}$) are set. These values are drawn from an uniform distribution in the interval $[0, 1]$ and are not modified during its lifetime.

**Setting cell fate.** When determining its fate at the next frame, each cell takes into account its $t_{div}$ and $t_{death}$, as well as the division and death curves of the current treatment. Specifically, the cell fate is set to division when the division curve's cumulative distribution function evaluated at the expected cell age at $f_{i+1}$ is greater than the cell's $t_{div}$. Similar reasoning follows for $t_{death}$ and setting cell fate to death.

$t_{div}$ and $t_{death}$ are defined in the $[0, 1]$ interval and have no temporal component, effectively acting as cumulative distribution functions whose derivative is the probability density function of the division/death curve.

In the simulation, the process of setting the cell fate is roughly equivalent to the following pseudocode:

```
next_cell_age = cell.age + delta_t  # cell will have this age at the next frame

if division_curve.cdf(next_cell_age) > cell.t_div:
    cell.fate = 'division'

else if death_curve.cdf(next_cell_age) > cell.t_death:
    cell.fate = 'death'

else:
    cell.fate = 'migration'
```

In its implementation of the pseudocode above, CloVarS randomly selects whether to first test for cell division or death, in order to avoid biases from always testing one before the other. Supplementary Fig. S1 presents three scenarios with numerical examples of setting the cell fate.

**Cell signals.** Each cell has an associated cell signal, which represents the expression level of a generic pathway, or the amount of an arbitrary molecule. The cell signal is a value in the $[-1, 1]$ interval that fluctuates over time, according to its type and parameters (Supplementary Fig. S3A). Currently, it allows simulations to test and compare fitness dynamics with the oscillations of an independent, individual value (Supplementary Video S2); the signal has otherwise no influence over the cell behavior or its fate.

When a cell division occurs, each daughter cell copies the value and type of the cell signal from their mother cell; afterwards, the signal from each cell fluctuates independently (Supplementary Fig. S3B). Supplementary Fig. S3C presents the currently implemented cell signal types.

#### Experimental data

**Cell culture.** All experiments were performed with A-172 glioblastoma cells (ATCC) on DMEM low glucose media (Gibco #31600-034) supplemented with 10% FBS (Laborclin #630111), 1% penicillin/streptomycin (Gibco #15140-122) and 0.1% amphotericin B (Gibco #15290-018). Cells were kept inside an incubator (Panasonic) at 37ºC and 5% $CO_2$ before and during the experiments. Cells were routinely tested for Mycoplasma contamination using the Mycoplasma Detection Kit MycoAlert (Lonza).

**Division times for the Control treatment.** For measuring the division times of cells under standard growth conditions (the "Control" treatment), cells were seeded in a 25 $cm^2$ cell culture flask (Corning) at low density (approx. 5000 cells). The flask was placed inside a Incucyte S3 live cell imaging system (Sartorius) for 96 hours. Brightfield images of 16 fields were captured every 30 minutes. After image acquisition, cells were manually tracked using ImageJ. Cell age at its division event was obtained by measuring the time interval between the mother cell mitosis and its daughter cells mitosis. Cells that escaped the field of view, did not divide, or did not complete a full cell cycle were not analysed. In total, 62 cell division events were evaluated.

**Division and death times for the TMZ treatment.** For measuring the division and death times of cells under Temozolomide (TMZ) treatment, cells were seeded in a 25 $cm^2$ cell culture flask (Corning) at low density (approx. 5000 cells). The flask was placed under a CytoSMART live cell imaging system (CytoSMART Technologies) for 15 days. At day 4, the culture media was refreshed and 50 µM TMZ (Sigma-Aldrich) was added. The culture media was again refreshed at day 7, removing the TMZ. Brightfield images of the field of view were captured every 5 minutes. After image acquisition, cells were manually tracked using ImageJ. Cell age at its division or death event was obtained by measuring the time interval between the mother cell mitosis and its daughter cells mitosis or death. Cells that escaped the field of view, did not divide/die, or did not complete a full cell cycle were not analysed. In total, 225 cell division events and 69 cell death events were evaluated.

#### Figure data

Code for reproducing the data from figures is located in the CloVarS github repository (https://github.com/jfaccioni/clovars). Data was obtained using CloVarS version 0.2.1 (different versions are not guaranteed to produce the same results).

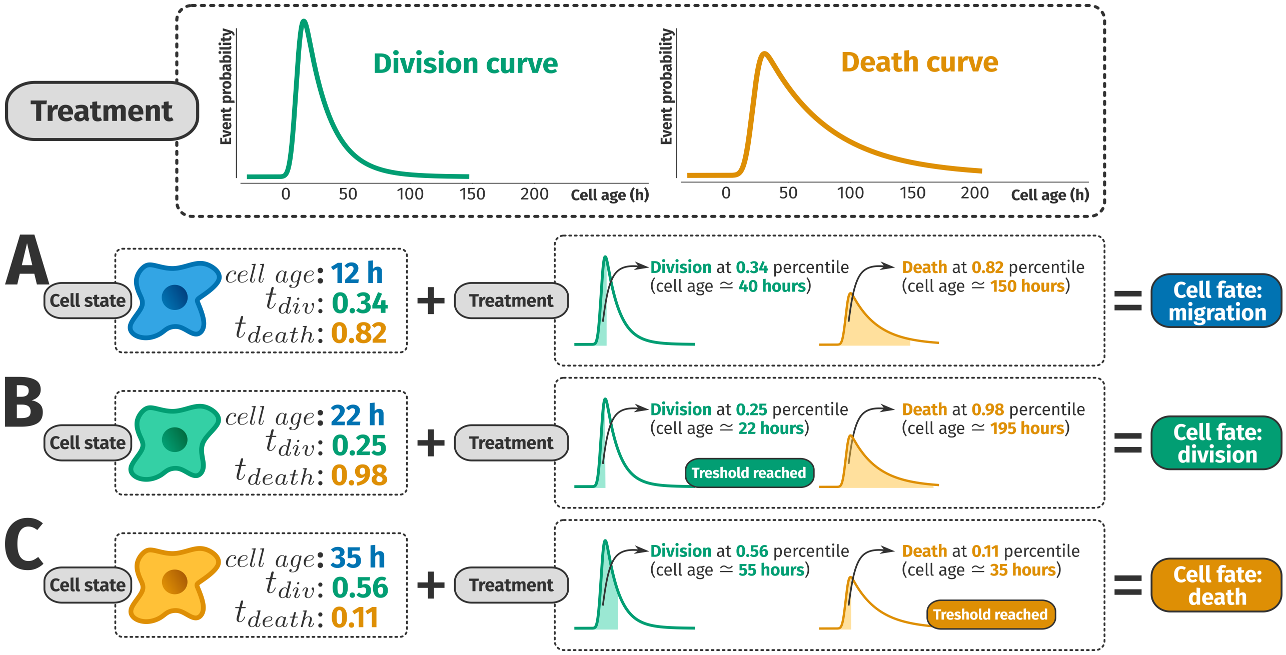


**Supplementary Fig. S1.** Numerical examples of cells whose fate at next frame is to (A) migrate, (B) divide, or (C) die.

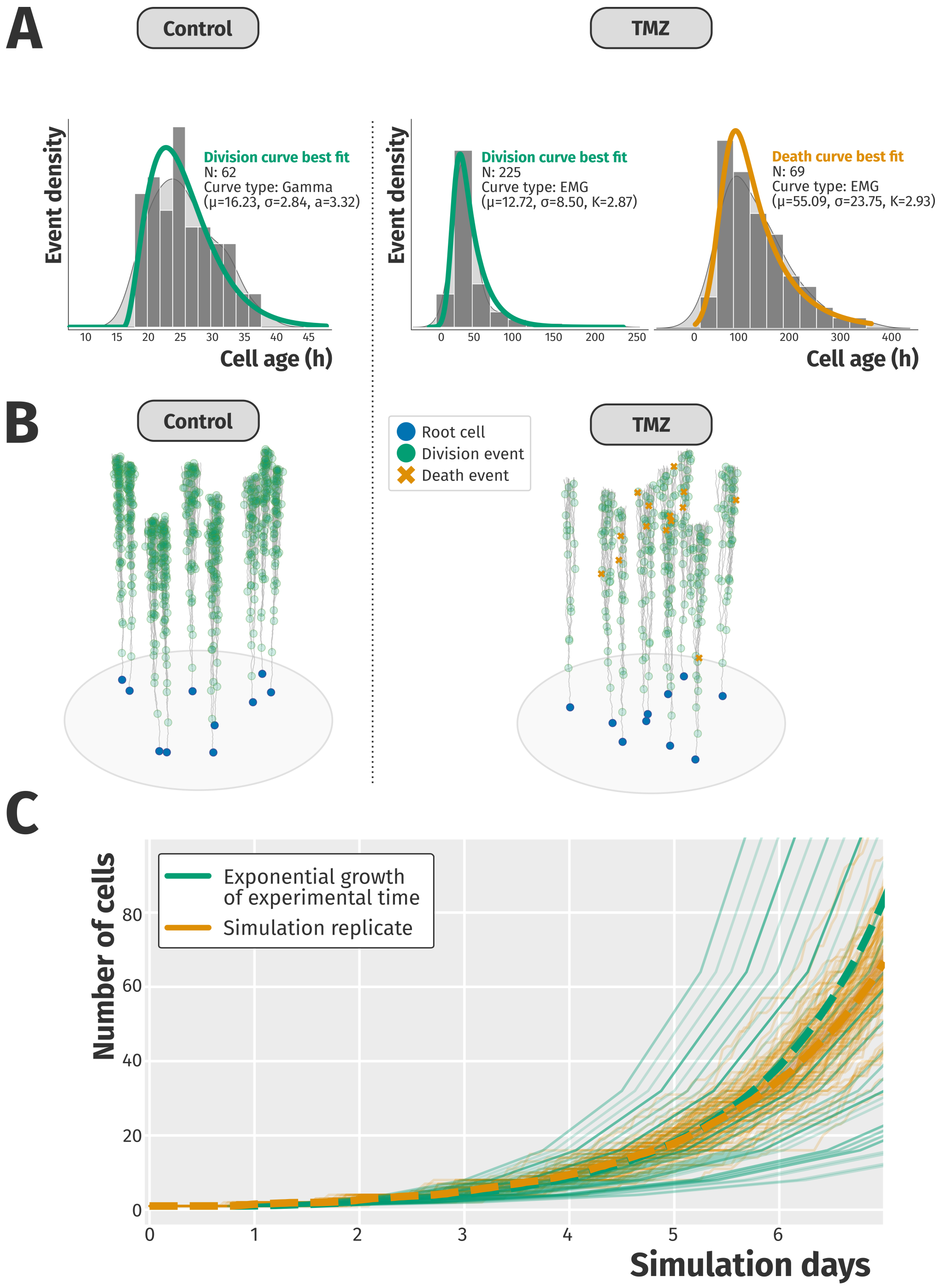


**Supplementary Fig. S2.** Deriving treatment parameters from experimental data. (A) Division/death curves were derived from experimental data (gray bars) of manually tracked single cell times to division/death, respectively. the gray curve represents a kernel density estimate of the experimental data. (B) 3D view of simulation results. The Z-axis corresponds to simulation time. Left: cells kept under the Control treatment for 7 days. Right: cells were under the Control treatment for 72 frames (3 days), then switched to the TMZ treatment. (C) Comparison between exponential growth using the division times in the Control treatment and 100 simulation replicates. The colony growth in the simulation closely resembles the expected growth rate of the experimental data. Dashed lines: replicate average. Code used to generate Supplementary Figs. S2A, S2B, and S2C can be found in the CloVarS repository.

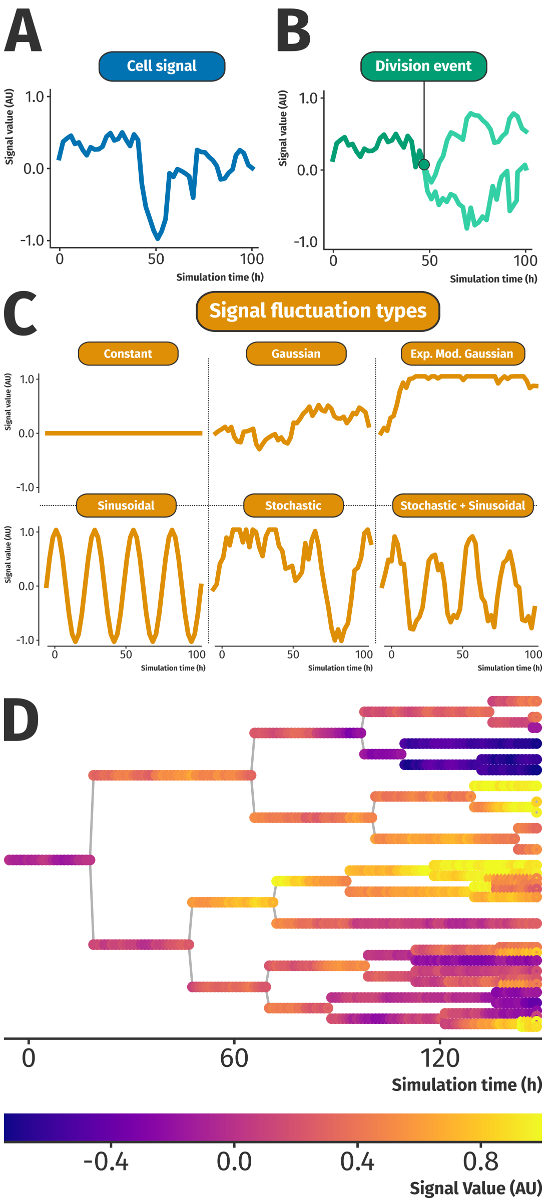


**Supplementary Fig. S3.** Cell signals in CloVarS. (A) During the course of the simulation, the cell signal oscillates between frames, fluctuating its value according to the fluctuation parameters. (B) Daughter cells inherit the cell signal value and fluctuation type from their mother, and oscillate it independently from one another. (C) Fluctuation types available for cell signals in CloVarS. (D) Signal fluctuation in an example lineage tree. A single colony was simulated for 144 frames ($\Delta t = 1h$). Signal type was Gaussian, with $\mu = 0$ and $\sigma = 0.05$.

**Supplementary Table S1. Parameters required to run CloVarS.**

| Name | Data Type | Description |
|---|---|---|
| Output folder | Text | Path to folder in which to save CloVarS output |
| Delta | Integer | Time (in seconds) between two adjacent frames |
| Frame to stop | Integer | Simulation ends when this frame is reached |
| Colony size to stop (single) | Integer | Simulation ends when at least one colony reaches this number of cells |
| Colony size to stop (all) | Integer | Simulation ends when all existing colonies reach this number of cells |
| Colony data | Structured | List of colonies to simulate (see Supplementary Table S2) |

**Supplementary Table S2. Parameters used to define a colony.**

| Name | Data Type | Description |
|---|---|---|
| Initial size | Integer | Initial number of cells in the colony |
| Radius | Integer | Cell radius (in µm) |
| Max speed | Integer | Cell max speed (in µm/s) |
| Fitness memory | Float | Value between 0 and 1 that defines how much of the mother's fitness is inherited by the daughter cells |
| Treatment regimen | Structured | Describes the treatment that the colony is subject to (see Supplementary Table S3) |
| Signal | Structured | Describes the signal dynamics of each cell in the colony (see Supplementary Table S4) |

**Supplementary Table S3. Parameters used to define a treatment.**

| Name | Data Type | Description |
|---|---|---|
| Name | Text | A name that identifies the treatment |
| Added on frame | Integer | The frame in which this treatment is added to the colony |
| Division curve name | Text | Type of curve to use for the treatment division curve (valid types: Gaussian, EMGaussian, Gamma, Lognormal) |
| Division curve parameters | Float | Parameters describing the shape of the division curve (mean, std, etc) |
| Death curve name | Text | Type of curve to use for the treatment death curve (valid types: Gaussian, EMGaussian, Gamma, Lognormal) |
| Death curve parameters | Float | Parameters describing the shape of the death curve (mean, std, etc) |
| Fitness memory disturbance | Float | Optional value for the colony's fitness memory after treatment is added to it |
| Signal disturbance | Structured | Optional type of signal dynamics for the cells in the colony after treatment is added |

**Supplementary Table S4. Parameters used to define a signal.**

| Name | Data Type | Description |
|---|---|---|
| Name | Text | Type of signal used (valid types: Constant, Stochastic, Sinusoidal, Stochastic + Sinusoidal, Gaussian, EMGaussian) |
| Initial value | Float | The signal's starting value (ranges from -1 to 1) |
| Signal parameters | Float | Parameters describing the nature of the signal's fluctuation (mean, std, etc) |

**Supplementary Movie S1.** Relationship between fitness dynamics and colony size. For each fitness memory ($f_m$), 100 colonies were simulated. The size (top) and mean $t_{div}$ (bottom) of each colony was plotted at each simulation frame ($\Delta t = 1$ h). As the colonies with low $f_m$ increase their size, their mean $t_{div}$ tend to stabilize around $0.5$ (i.e. the mean value of the uniform distribution in the $[0, 1]$ interval). In contrast, colonies with greater $f_m$ tend to propagate the $t_{div}$ of the initial cell across generations. This results in greater colony size variability for larger $f_m$ values. The code used to generate the Supplementary Video S1 can be found in the CloVarS repository.

**Supplementary Movie S2.** Example of signal fluctuation dynamics. For each fitness memory ($f_m$), 100 colonies with a Gaussian signal starting at $0.0$ were simulated. The signal mean (top) and variance (bottom) for each colony was plotted at each simulation frame ($\Delta t = 1$ h). CloVarS allows researchers to explore mitotic-independent cell signal dynamics as colonies grow in size. The code used to generate the Supplementary Video S2 can be found in the CloVarS repository.
